# Protein Family Classification using Deep Learning

**DOI:** 10.1101/414128

**Authors:** K S Naveenkumar, Babu R Mohammed Harun, R Vinayakumar, KP Soman

## Abstract

Protein classification is responsible for the biological sequence, we came up with an idea which deals with the classification of proteomics using deep learning algorithm. This algorithm focuses mainly to classify sequences of protein-vector which is used for the representation of proteomics. Selection of the type protein representation is challenging based on which output in terms of accuracy is depended on, The protein representation used here is n-gram i.e. 3-gram and Keras embedding used for biological sequences like protein. In this paper we are working on the Protein classification to show the strength and representation of biological sequence of the proteins.

## 1. Introduction

Human body comprises of the many cells the key for the formation of all these are the DNA(Deoxyribonucleic acid) which is a chain of nucleotides that are thread like chain in nature responsible for carrying genetic instruction for development, functioning of organisms in the body and RNA(Ribonucleic acid) which is a polymeric molecule which is very much essential for the three biological roles namely CDRE(Coding, Decoding, Regulation and Expression) of each and every gene present in the body, which are present in every living being. Like human beings use various language for communication, biological organisms use these type of the codes by using DNA and RNA for the communication. Selection of the type of feature extraction is a challenging task because it helps for the studying of the types of the genes to the machine using the machine learning algorithm, Even a most highly sophisticated algorithm would go wrong if the feature extraction is not done in a proper form. The features from the existing data can be obtained by manually or by using unsupervised (data without labels) fashion[1],[2]. This work focuses on protein family classification wit the publically avilable data Swiss-Prot[3]. In this work application of keras embedding and n-gram technique is used to map the protein sequences into numeric vectors and followed by traditional machine learning and deep neural network (DNN) for classification.

The rest of the paper are organized as follows. Section 2 discusses the related works in a detailed manner. Section 3 presents the background works. Section 4 presents the description of the data set. Section 5 gives a overview about the proposed architecture which is used in building this work. Section 6 and 7 presents results and conclusion, future work directions and discussions.

## 2. Related works

In this paper we mainly focus on the related works done previously on protein classification using deep learning. Using Natural language processing which mainly focus on biological sequence, the features are been extracted using n-grams, Bio-vec and Prot-v to identify the protein structure and classified into different structure for which deep neural network has been implemented to find n-dimensional vectors[4][5]. These are considered as main features, using some of CBOW and skip-gram methods, word2vec, DM and DBOW for doc vec consider another method in which we can classify the protein biological embedding sequences[1].In which supervised learning has been used as classifier to classify the predictive performance of the data, from the data that has been classified various features were been extracted and compared, after which the accuracy based classification is done[6]. Using the extracted features and considering the values, prediction is done by using Random forestry, Naviers bayes classifier and Decision tree methods. The outputs of these are taken under consideration, which is helpful in prediction solving of protein classifications and time efficient[7][8]. In proteins the transformation of sequence of cells requires an analysis to obtain source function of proteins to obtain this sequence gaussian process regression is generalize to find number of variables[9]. By training the sequential data of various proteins regarding the set function analysis accuracy and result can be obtained [4]. The proteins are classified based on the sequence with a distribution of the ten classes based on the algorithm which has been developed recently which is an advanced level of the deep learning that is by the means of the ELM (Extreme machine learning). When this algorithm is tested by comparing it with the old that is with the existing algorithm which is none other than the BNN (Backpropagation neural network) method it had a significantly greater response when compared with the BNN, the ELM requires very less time and four times magnitude less training when compared with BNN with a greater accuracy. The ELM (Extreme machine learning) which is used in this architecture of network do not have any of the parameters for controlling automatically only manual operation is needed which is easy to be tuned to the machine[10]. The classification are the implementation part of the data into the algorithm some of the types of the classifiers that are being implemented are GE (Genetic algorithm), Fuzzy ARTMAP which are being used in the deep learning for classification that has to improve the accuracy. This deals with the GE, Fuzzy ARTMAP and RCM (Rough set classifier model)[11].The classification of the protein is a tough task for the persons to identify their family because it is tedious time consuming process now a days the machines are being used for the classification based on the machine learning techniques such as the SVM(Support vector machine),RF(Random forest).K-NN(K-nearest neighbor) and Fuzzy K-NN are being used[12].

## 3. Background

### 3.1. Proteomics

Proteomics is the study of proteins which is concerned with the protein expression analysis of a cell or an organism. Proteins are molecules that contains one or more long chains of amino acids which performs variety of functions such as catalytic metabolic reaction, DNA replication, Responding to stimuli and transporting molecules inside the cell. The amino acids in the proteins are formed of sequences which are stated by their genes of the nucleotide which results in protein folding into a three dimensional specific function. Proteome can be expressed as the inverse of the entire protein which can be shown by an organism, tissue or by a cell. They vary with respect to time and their requirements based on the stresses of the cells[4]. Proteomics which are combination of various branches has got contributed from genetic information of HGP (Human Genome Project) which is a blooming topic under research of proteins. Proteomics shows that it is responsible for the large scale experiment analysis of proteins and is specifically used on protein purification and mass spectrometry[1].

### 3.2. Genomics

Genomics a branch of the molecular biology which is a study related to the function, evolution, structure, editing and mapping of genomes. Genome consists of the haploid (single set of unpaired chromosomes) set of chromosomes in each cell of multicellular organism or in gametes of microorganisms they are also referred to a complete set of genetic material or genes present in a cell or organism. An organisms complete set of DNA including all its genes is Genome[6]. On the other hand Genetics means study of genes which are individual and their functions in inheritance, whereas genomics is the study of characterization which are collective and genes of qualification, that recalls the production of the proteins in a method that is direct that is with the AOE (assistance of enzymes) and MM(messenger molecules).

### 3.3. N-gram

Humans used to communicate with each other for the purpose of the communication in the same way the computers communicate with its users by the means of the algorithm that is by performing the task which was assigned by the coder to the machine. The programmer uses the certain algorithm to ask the computer rather the machine to perform certain task. As years passed by the computers are taught linguistics in the form of the data put in an algorithm with the help of the probability, as a result a new field is emerged that is computational linguistics along with probability. Many algorithms are being created for the feature extraction from the data, out of which n-gram is one of the feature extraction which is being followed majorly by all. N-Gram is a n items of the given sample which has a sequence of the text or speech those can be of the syllables, words, corpus (which is used for the text data), or text. Based on the type of the words, letters or a text that are used they are classified as “unigram” means taking one word or a letter or a text then as the “bigram” taking two at a time and so on such as “three-gram”, “four-gram”, “five-gram” etc. They are extensively used in the NLP(Natural Language Processing) and Speech Recognition which is a statistical and used for the extraction of the phonemes(distinct sound unit related to a particular language that are separated)and sequence of phonemes.

### 3.4. Keras embedding

Word embedding deals with the vector representation of the dense vectors. This method is much more accurate than that of the previous methods that was used which is the bag of words. Keras is the library function which allows the use of the embedding for the word semantically and syntactically. This is an API data as an input which can be prepared by the Tokenizer which has to be integer encoded. They are initialized with the random weights that are responsible for the embedding in the training set.

In recent days, Keras embedding has performed well in compared to n-gram as text representation method in most of the applications[13],[14],[15],[16],[17],[18].

### 3.5. Machine learning

Naives Bayes The naives Bayes is a classifier in the machine learning which uses the probabilistic methods alongside with the help of the bayes theorem, which is easily scalable with linear number of parameters such as the features or the predictors that help the machine to learn about the problem, is very efficient method of classifiers which has a very greater level of accuracy that are easy to learn, which mainly focus on the dependent attributes and they can easily deal with the parametric attributes which are continuous. The process of maximum training takes place by the method of the closed form evaluation expressions that takes in a linear time for the completion than that of the other method that is the iterative process of approximation. This algorithm has a technique that has the capability of assigning the labels that are given to the class with the problem instance which are represented in the form of the vectors of the values that are featured and the values of the class are taken from the set that is finite. There are two types of the method in the machine learning that is the supervised and the unsupervised machine learning this algorithm goes well with the supervised machine learning and has high efficiency on the machine learning. This has a greater advantages for the users who are using this algorithm for the various purposes is that this need a very little amount the training data set when it is being compared with the other types of the algorithms and gives a greater accuracy.

#### 3.5.1. Decision tree

A method which can be said as an algorithm that only depend on the statements which are conditional and control, uses the tool which takes the decision that has the tools which supports or agree with the graph which are like the tree with the models that have decision which can have a possible outcomes that also include the outcomes of the event chances along with the utility and the costs of the resources are the decision trees. This is a machine learning tool that is so popular which are used for the research operations because this type of the algorithm deals with the various types of the analysis mainly decision analysis that helps in identifying a problem or a situation which are similar in the case of the reaching a goal. The decision trees are in the structure of the flowchart which has nodes which are internal and they are referred by the term which is coined to be as “Test”, that has a representation of the of the results of the test in which it has a leaf node and each of the leaf node represents the class to which the label belongs. In the analysis done by the decision tree which can be also referred as the DA (decision analysis) they are form a close resemblance to that of the influence diagram which are used for the various other methods such as the VAD (visual, analytic and decision support method) by which the values of the alternatives which are competing can be calculated.

#### 3.5.2. Random forest

Data is a point in space which has an infinite dimension, this century is the one which deals with data than with people large amount of data are found and they are increasing day by day so as the machine learning algorithms are used for the studying and dealing with data, one of the algorithm which deals with the data better are the random forest tree method which are used for solving the huge data sets. Algorithms of the recent trend are focused on the accuracy of the data even though they are focused on the accuracy they fail to classify the data with greater accuracy. So this algorithm that is random forest tree is used for the classification of the large amount of the data with the regression technique. Machine learning techniques are the recent trend in this era so this is used along with the random forest tree technique as an ensemble this leads to a cutting edge technique in the field of the data mining. Ensemble is the process of the data mining that consists of many classifiers who are individual for the classification of the data and to create a new data of instances. Random forest technique is one of the most famous and well know technique for ensemble classification because it has a lot of features present in it. Random forest trees are formed by the combination of predictors of the trees such as each of them depend upon the value of sampled random vector which are independent of each other with the similar type of the distribution to all the types of the trees that are present in that forest. The errors that occurs in this type of the trees are much reduced as the number of the tress that are formed are increased and become large. They are generalized so that the occurrence of the error is reduced even if the error occurs they depend on the strength of the trees that are individual, that belong to that forest and the relation between them. By using this method the features are splitted so that error rates are decreased and completely favorable with reduced noise.

### 3.5.3. AdaBoost

This method is not new to machine learning by the means of the name this may find different but the expansion for the ADABOOST is Adaptive Boosting, used by the weak learners for the coding kidding, they are also called as the discrete adaboost because it follows the classification method in the process when compared to regression. This type of the algorithm is used for boosting the performance of the machine learning as the name suggest. The performance of this type of the algorithm is for boosting the performance on the decision trees on the classification which are binary none other than the binary classification problem. The algorithm which is most suited with this type are the decision trees so they are used along with the adaboost algorithm, since these trees are short they contain only one decision tree for the purpose of the classification so they are called as the decision stumps. The weight is weighed at each instance of the dataset at the training level, the initial level of the weight is set to the weight that is given by the formula given below

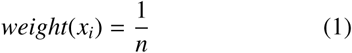

where *x*_*i*_ is the training instance at the *i* the level and *n* is the training instance.

AdaBoost classifiers are a meta-estimator which begins by fitting classifiers on the datasets that are original, they are additional to that of the classifier which are original. The weights are incorrectly distributed at any instances that are adjusted to classifiers which are subsequent and focuses on difficult tasks much more.

#### 3.5.4. Support Vector Machine(SVM)

Support Vector Machine(SVM) is the classifier in the machine learning which stands for the Support Vector Machine (SVM) which means formally as the discriminative classifier which are separated by the hyperplane. There are three methods of the learning in the machine learning they are the supervised and unsupervised and reinforcement learning the svm deals with the supervised machine learning which means that it consist of the dependent variable that can be predicted using the predictors which is nothing but the independent variables. Here the support vector machine that is the SVM follows the supervised learning (which deals with the datasets that are labelled) along with the other algorithms that help in the analysis of the data that are used for the regression and the classification analysis. They are successful in the high dimensional spaces and also were the number of the dimensions are greater than the number of the samples. It uses the decision functions which are called as the support vectors for the purpose of the training the points in the subset so that the memory would be efficient.

#### 3.5.5. k-nearest neighbors(KNN)

The k-nearest neighbors(KNN) stands for the K nearest neighbor algorithm which is a non-parametric methods which are used in the classification of the data and in the process of regression. In the both the methods which are stated above the input factors consists of the k closest examples for the training in the feature space, but the output depends upon the k-NN whether they are used for the process of the classification or for the regression. In this method the training data are the vectors that are in the multidimensional space which are in the feature with the each class with the label. During the training of the algorithm it consists of only the storing of the vectors which belong to the feature called as the feature vectors and the class to which the labels belong of that of the training data. When the k-NN is used in the classification the output which is got after the application of the algorithm is calculated as the class which is having the highest frequency from the base of the K-most similar instances, each of the instance are responsible for the votes of the class which are taken as the prediction.

### 3.6. Deep learning

Deep learning is a sub model of machine learning which has the capability to obtain optimal feature representation from raw input samples[19]. Deep neural networks (DNN) is a method of the machine learning tools, which allows one to learn about the complex functions which are non-linear from a given input which reduces the error cost. Recurrent Neural Networks (RNN) is a method of the machine learning tools, which uses the sequential data since each neuron has the internal memory in them to store the information about the input which had occurred already[19]. RNN generates vanishing error gradient issues when try to learn the long-term temporal dependencies. Long short-term memory commonly referred as LSTM (or blocks) are the main units in the building of layers of a RNN (Recurrent Neural Network). This has the capability to handle vanishing and error gradient issue. LSTM has a memory which has the gates they are the input gate, output gate and the forget gate and a self-recurrent connections with that of the fixed weights. The cell which is present is the main responsible for the memory.

### 3.7. Protein Space Construction

Set of proteins which are evolutionarily related, similarly involving in structures or functions belongs to protein family. The protein family consists of the higher classifier named G-protein coupled receptors, based on which they are divided given in the picture below. The protein family is represented as the ‘Pfam’ which are classified as the “Family” and “Domain”[4]. ‘Pfam’ which belongs to the protein classification which are made by the biologist for the computational purpose includes a view of description of the family, multiple alignments, protein domain architectures, examine species distribution. Classification is the task next to feature extraction which determines the value for the data set that has been got. In this paper the classifier used for classifying protein family is SVM (Support Vector Machine) based on the sequences that is on primary structures this classifier classifies the given data. SVM (Support vector machine) are the supervised learning methods which are used by machine learning for analysis such as data and regression based classification[6].

### 3.8. Protein Space Construction

The main objective of this process is to create a sequence in biology which is a distributed representation. The data is divided into two they are training data and the test data, the system is trained by the means of the training data and with the help of the test data it is verified. For this many process such as the ten cross ten is used to check the train data with the test, many algorithm are being used for this process, based on the accuracy which is being got and to improve the accuracy the algorithm is varied[4]. A large set of the training data is required for the machine to learn and based on the information present in the data, here it is the protein sequence machine learns as we learn and the output is when a sequence is being given to the machine rather the algorithm it tells to which sequence it belongs to easily within no time. The n-gram algorithm uses the information from the protein that are used for the lapping of the window from three to six residues. Using a two cross fold cross validation results are more accurate for different window size.

### 3.9. Protein Space Analysis

The protein space analysis is used for many process that are used in training space for the analysis of the spread over of the physical and chemical biological properties that are used in the n-gram more specifically the three gram is used in the hundred dimensional space to the two dimensional space by the means of the SNE (Stochastic Neighbor Embedding). Some properties like the volume and mass were being taken into consideration for the classification of the data, the protein space characteristics have been studied and from Lipschitz constant[4]

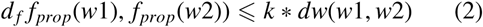

Were f, is scale of the properties on which the n gram has been used, D, is the metric distance. Df, is the score differences of the absolute value. Dw, is the distance that is Euclidian between the 2 three-grams w1 and w2

## 4. Description of data set

We gathered family information of about 40433 protein sequences in Swiss-Prot from Protein family database(Pfam), which consists of 200 distinct families. Swiss-Prot is a curated database of primary protein sequences which is manually annotated and reviewed. There is no redundancy of protein sequences in the database and is evaluated based on results obtained through experiments. The data contain 84753 protein sequences[3].

## 5. Proposed Architecture

An overview of proposed architecture is shown in Fig 1. This contains a protein sequences. These sequences are passed into protein representation layers. This layer transforms protein into numeric vectors i.,e. dense vector representation. There are two types of representation are used. They are n=gram and Keras embedding. In n-gram we used 3 gram with feature hashing of length 1000. These vectors are passed into different deep learning layers such as dnn, rnn, lstm and CNN for optimal feature extraction. DNN model contains 5 hidden layer each hidden layer contain 128 hidden units. RNN and LSTM layer contains 128 units and memory blocks respectively. CNN contain 128 filters with filter length 4. CNN layer followed pooling layer. We used max-pooling which is of length 3. These feature are passed into fully connected layers for classification. The fully connected layer contains connection to every other neurons. This used softmax as an activation function. To reduce the loss, categorical cross entropy is used. The categorical cross entropy is defined as follows.

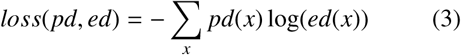

where is true probability distribution, is predicted probability distribution. We have used as an optimizer to minimize the loss of binary-cross entropy and categorical-cross entropy.

## 6. Results

All experiments of deep learning algorithms are run on GPU enabled machines. All deep learning algorithms are implemented using TensorFlow[20] with Keras[21] framework. The 5-fold cross validation accuracy of each deep learning models is reported in Table 1. The hyper parameters for DNN, RNN and LSTM is done through following hyper parameter selection methods. 3 trials of experiments are run for unit. Memory blocks varying in the range 32,64,128 and 256. Experiments with 128 performed well. Thus we decided to set 128. All experiments are run for learning rate in the varying range 0.01 to 0.5. Experiments with learning rate 0.02 preformed well. For rnn and LSTM, when we increase the number of hidden layer to more than one the performance is reduced due to over fitting. Thus we decisded to use only hidden layer for RN and LSTM networks. For DNN layer 2 trails of experiments are run with 1, 2, 3, 4 and 5 layer. Experiments with 5 layer performed well. All these experiments are run till 100 epochs. The best performed model again run for 1000 epochs and the detailed 5-fold cross validation accuracy of various deep learning models are reported in Table 1.

**Table 1:** Results of deep learning model with n-gram and Keras embedding

| Model | cross validation |
| --- | --- |
| DNN with 3 gram | 81.2 |
| RNN Keras embedding | 88.41 |
| LSTM Keras embedding | 91.24 |
| CNN Keras embedding | 90.18 |

## 7. Conclusion, Future work and Discussion

This paper has proposed deep learning method for protein classification. To transform protein to numeric vector, the n-gram and Keras embedding representation is used. Deep learning method with Keras embedding has performed well in comparison to the n-gram with deep neural network. The main reason is due to the Keras embedding has the capability to preserver the sequential information among the protein sequences. Thus, the deep learning algorithms able to capture the optimal information from the syntactic and semantic information of Keras embedding vectors. The proposed methodology can be employed for other domain in biology tasks such as genomics, DNA classification. This is one of the significant directions towards future work.

